# REVA: a rank-based multi-dimensional measure of correlation

**DOI:** 10.1101/330498

**Authors:** Bahman Afsari, Alexander Favorov, Elana J. Fertig, Leslie Cope

## Abstract

The *neighbors* principle implicit in any machine learning algorithm says that samples with similar labels should be close to one another in feature space as well. For example, while tumors are heterogeneous, tumors that have similar genomics profiles can also be expected to have similar responses to a specific therapy. Simple correlation coefficients provide an effective way to determine whether this principle holds when features and labels are both scalar, but not when either is multivariate. A new class of generalized correlation coefficients based on inter-point distances addresses this need and is called “distance correlation”. There is only one rank-based distance correlation test available to date, and it is asymmetric in the samples, requiring that one sample be distinguished as a fixed point of reference. Therefore, we introduce a novel, nonparametric statistic, REVA, inspired by the Kendall rank correlation coefficient. We use U-statistic theory to derive the asymptotic distribution of the new correlation coefficient, developing additional large and finite sample properties along the way. To establish the admissibility of the REVA statistic, and explore the utility and limitations of our model, we compared it to the most widely used distance based correlation coefficient in a range of simulated conditions, demonstrating that REVA does not depend on an assumption of linearity, and is robust to high levels of noise, high dimensions, and the presence of outliers. We also present an application to real data, applying REVA to determine whether cancer cells with similar genetic profiles also respond similarly to a targeted therapeutic.

**Author summary:** Sometimes a simple question arises: how does the distance between two samples in multivariate space compare to another scalar value associated with each sample. Here, we propose theory for a nonparametric test to statistically test this association. This test is independent of the scale of the scalar data, and thus generalizable to any comparison of samples with both high-dimensional data and a scalar. We apply the resulting statistic, REVA, to problems in cancer biology motivated by the model that cancer cells with more similar gene expression profiles to one another can be expected to have a more similar response to therapy.

## Introduction

The venerable K-nearest-neighbor approach succinctly captures a principle at the heart of any machine learning algorithm: samples with similar labels should be close to one another in feature space as well. Here, we present a method for quantifying the extent to which the neighbors principle holds in a given dataset, without making specific model assumptions. Our method, called REVA, was motivated by computational genomics for cancer, where machine learning methods are applied to high dimensional genetic signatures to identify aggressive tumors and predict response to therapy. The biology behind response to therapy is complex, with extensive heterogeneity of genetic profiles and therapeutic response between distinct tumors and even within the cells that comprise a single tumor. Yet, the neighbors principle can be expected to hold when predicting therapeutic response of targeted therapies that work by blocking a genetic alteration specific to a tumor. Namely, tumors that are similarly responsive to that therapy are hypothesized to have more similar genetic profiles than tumors do not, reflecting the wide variety of mechanisms that individual tumors can utilize to escape treatment by targeted therapies. It is of interest, then, to have computationally efficient, model-free statistical measures of the extent to which samples with similar profiles share similar responses.

To develop the REVA method to quantify the extent to which the neighbors principal holds, we adopt the following notation. For each sample *i*, we observe (**x**_*i*_, *y*_*i*_), where *y*_*i*_ is a scalar, response variable and **x**_*i*_ is a vector of predictors. In our example of targeted therapeutic response in cancer, *y*_*i*_ would be a measurement of therapeutic response and **x**_*i*_ a genetic profile for tumor *i*. According to the neighbors principal described, we expect small values of |*y_i_*-*y*_*j*_| to correspond to small values of *D*(**x**_*i*_, **x**_*j*_), where *D* is a distance on **X**. Our goal is then to measure the correspondence, assign confidence intervals, and perform hypothesis tests. The resulting measure should capture both linear and non-linear relationships, be relatively invariant to the dimensionality of the **X**, and statistical procedures, including hypothesis testing, should be computationally efficient. The cancer genomics context of our motivating example suggests some additional constraints on the possible solutions to the problem. Specifically, genomics data is subject to pervasive, technology-specific biases which can be controlled using rank-based analysis procedures as shown in the literature ([1–4]).

Recent work has led to the development of a small but growing class of generalized correlation coefficients applicable to multidimensional data. These methods were pioneered by Szekely, Rizzo and Bakirov with their development of *distance correlation*, wherein interpoint distances *D*(**x**_*i*_, **x**_*j*_) and *D*′ (**y**_*i*_, **y**_*j*_) are calculated for all pairs of vectors (**x**_*i*_, **x**_*j*_) and (**y**_*i*_, **y**_*j*_) and then based on these calculations, Pearson’s correlation is calculated as *ρ*(*D*(**x**_*i*_, **x**_*j*_), *D*′ (**y**_*i*_, **y**_*j*_)) [5–7]. Heller, Heller and Gorfine [8] presented an elegant alternative, calculating a rank-based correlation coefficient, but requiring that one sample be chosen as a reference point and ranking the other samples according to their relative distance from the selected reference. More recently, Shen, Priebe, Maggioni and Vogelstein [9] extended the distance correlation framework of Szekely and colleagues to restrict the correlation to specific scales relevant to the data, rather than weighing all pairwise relationships among variables equally [9]. None of these methods depends on parametric assumptions, all are similarly computationally efficient and all three have the potential to capture a variety of linear and non-linear relationships. However, only the approach by Heller *et al.* [8] is based on ranks, and it requires to choose one of the samples as the reference point, and the result depends on the choice. Using a very different, generalized approach to the same problem, Gretton, Fukumizu, Teo et al. introduced the Hilbert-Schmidt independence Criterion, a kernel dependence test in multidimensional Euclidean spaces [10–12]), building on earlier kernel methods like N-distances ([13]).

REVA was inspired by the Kendall rank correlation coefficient. Briefly, Kendall’s statistic starts by designating a pair of samples (*x*_*i*_, *y*_*i*_) and (*x*_*j*_, *y*_*j*_) as concordant if *x*_*i*_ *> x*_*j*_ when *y*_*i*_ *> y*_*j*_, and discordant if the order is not the same, and is then defined as

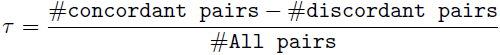

We observed that if we considered triplets of samples rather than pairs, then it was possible to define concordant and discordant states, based on the relative order of the pairwise distances, regardless of the dimensionality of the data, without the requirement that one sample be selected as a reference point. This work is a natural extension of our previous work using Kendall’s statistic to compare genetic dysregulation between tumors of two subtypes or tumors relative to normals [14, 15] using analytical hypothesis testing methods developed in [16], building upon previous permutation tests developed for similar analyses [17]. In the following sections we develop the REVA statistic, using the theory of U-statistics to demonstrate consistency and asymptotic normality. To establish the admissibility of the REVA statistic, and explore the utility and limitation of our model, we compared it to the most widely used distance based correlation coefficient in a few simulated conditions, and demonstrated it in a real application associated genomics profiles in cancer cells with targeted therapeutic response.

## The REVA Statistic

Like Kendall’s *τ*, REVA starts with a definition of concordance. Consider any 3 samples, **x**_*i*_, *y*_*i*_, **x**_*j*_, *y*_*j*_, and **x**_*k*_, *y*_*k*_ where the **x** are vectors and the *y*, are scalar and suppose, without loss of generality, that *y*_*i*_ *< y*_*j*_ *< y*_*k*_ so that the jth sample represents the median of these 3 points. Inspired by Frechet’s generalization of the median, we borrow the notion of the median as a point whose distance to other points is minimum on average, rewriting the necessary condition as follows, *y*_*j*_ is the median if |*y*_*i*_-*y*_*k*_ |*> max*(|*y*_*i*_*-y*_*j*_ |, |*y*_*j*_*-y*_*k*_). We will say that the triplet is concordant if **x**_*j*_ also represents the *Frechet median* among the *x*’s so that *D*(**x**_*i*_, **x**_*k*_) *> max*(*D*(**x**_*i*_, **x**_*j*_), *D*(**x**_*j*_, **x**_*k*_)). REVA is then defined as the proportion of all triplets that are concordant. This concept is formally developed in the following definitions.

### Definition 1

*Let D be any metric on space* **X**. *We define the median sample, out of three arbitrarily selected samples***x**_*i*_, **x**_*j*_, **x**_*k*_ *∈* **X** *as the one satisfying* **M**(**x**_*i*_, **x**_*j*_, **x**_*k*_) = arg min_*a∈*{*i,j,k*}_ *Σ*_*b∈i,j,k*_ *D*(**x**_*a*_, **x**_*b*_).

### Remark 1

*In the development of REVA, and in the theoretical results that follow, we generally assume that all* **x**_*i*_ *are unique, as well as all distances, D*(**x**_*i*_, **x**_*j*_) *so that there is a unique* **M** *for each triplet. In practice there may be ties, of course, in which case, we propose to break the tie randomly (e.g. selecting between two possible medians with a probability of 0.5 for each).*

### Remark 2

*Because we are using ranks of distances we are not calculating a median directly but only identifying the median sample* **M**(**x**_*i*_, **x**_*j*_, **x**_*k*_). *In the remainder of the paper we will use the terms median sample and median interchangeably to refer to the sample. To avoid confusion, we use* **M** *to indicate the median vector (i.e. in* **X** *space) and m to denote the scalar median sample (i.e. in ℛ).*

Now, we are ready to define REVA itself.

### Definition 2

*Let* **Z**_1_ = (**X**_1_, *Y*_1_), *…,* **Z**_*n*_ = (**X**_*n*_, *Y*_*n*_) *be i.i.d. samples where* **X**_*i*_*’s are random vectors ∈ X and Y*_*i*_*’s are their i.i.d. corresponding scalars (∈ R). A triplet, i, j, k, is concordant if* **M**(**X**_*i*_**X**_*j*_, **X**_*k*_) = *m*(*Y*_*i*_, *Y*_*j*_, *Y*_*k*_) *and discordant otherwise. We define REVA as the proportion of concordant triplets, expressed mathematically as,*

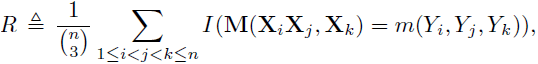

*where I*(*x*) *is the indicator function which is 1 if x is true and 0 otherwise.*

### Remark 3

*With the definition of median in hand for multivariate triplets, the REVA framework is readily extended to accommodate a vector-value Y. The criteria for concordance becomes* **M**(**X**_*i*_, **X**_*j*_, **X**_*k*_) = **M**(**Y**_*i*_, **Y**_*j*_, **Y**_*k*_), *and with this change, the definitions above accommodate the generalized scenario, and the asymptotic theory that follows holds with some changes to the variance calculation.*

The expected value of REVA (denoted by 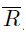) is easily derived under independence. It is the probability that **M**(**X**_*i*_**X**_*j*_, **X**_*k*_) = *m*(*Y*_*i*_, *Y*_*j*_, *Y*_*k*_) by chance alone. Trivially, if the *X*_*i*_ are independent of *Y*_*i*_ then all the possibilities are equally likely. Hence,

### Proposition 1

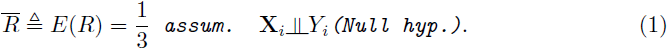

*Also if ∀ i, j, k, l D*(**X**_*i*_, **X**_*j*_) *> D*(**X**_*k*_, **X**_*l*_) ⇔ |*Y*_*i -*_ *Y*_*j*_*|> |Y*_*k -*_ *Y*_*l*_ *|, then 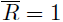 which means if there is perfect matching between the pairwise distance and the scalar, REVA captures it perfectly.*

### Remark 4

*It is easy to see that the neighbor principle holds, and 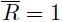 when* **X** *and Y are perfectly anticorrelated as well (D*(**X**_*i*_, **X**_*j*_) *> D*(**X**_*k*_, **X**_*l*_) *⇔ |Y*_*i*_ *Y*_*j*_*| < |Y*_*k -*_ *Y*_*l*_, *| for all i, j, k, l.) However 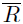 can take on values below*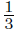 *in unusual cases where the order relationships among the d***X**_*i*_*s is very different from the Y*_*i*_*s. For example, 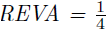 if x and Y are both scalar values with 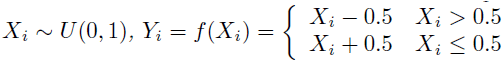*

## Asymptotic Normality and Implementation

In some settings, it will be desirable to calculate a confidence interval or perform a test and assign a p-value for these associations. Bootstraps and permutations provide a general method to establish a null distribution and calculate relevant statistics. However, this process can become computationally intensive especially if we need to correct for multiple hypothesis as with False Discovery Rate adjustment [18]. As is customary in statistics and machine learning, we attempt to find the asymptotic distribution for REVA and as usual we anticipate its asymptotic normality. U-statistic theory provides the theoretical framework for establishing the asymptotic normality of REVA:

### Theorem 1

*As the number samples (n) grows, REVA converges asymptotically to a normal distribution, i.e.*

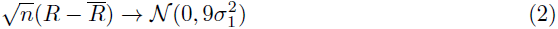

*where* 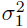 *is defined below.*

### Proof 1

*Since the indicator function is a bounded function, we can simply apply the main result of the U-Statistic theory [19] which proves the theorem.*

To calculate the variance, 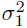, we define

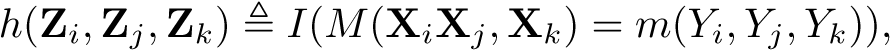

and 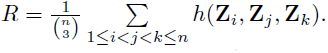.

The main theorem of U-Statistics says that for large samples, 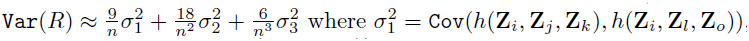, 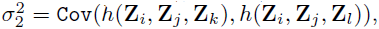 and 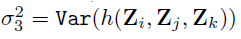. To calculate a p-value, it is necessary to estimate these parameters under the null hypothesis, i.e. when 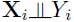. An expansion of 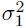 under that circumstance is shown below, similar expressions for 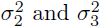 are not shown.

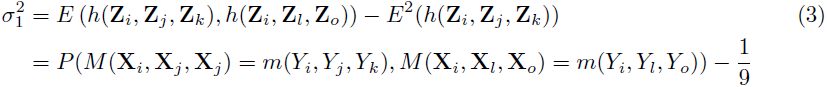

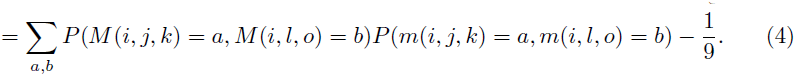

The second equality follows from the observation that *h* is an indicator function and the third equality from the law of total probability and independence of **X**_*i*_’s and *Y*_*i*_’s under the null hypothesis. Taking advantage of the symmetry between **Z**_*i*_, **Z**_*j*_, and **Z**_*k*_, we can show the full distribution of possible values for equation 5 in a simple tabular form (left side of Table 1). We note that because of the symmetry, it is necessary to estimate only one parameter: *α* = *P* (*M* (**X**_*i*_, **X**_*j*_, **X**_*k*_) = *i, M* (**X**_*i*_, **X**_*l*_, **X**_*o*_)) = *i*. In the absence of ties, we can pre-compute the exact values for scalar-valued, ranked data, as shown on right side of Table 1. Table 2 shows the similar probabilities that are obtained for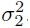. Once again the scalar matrix can be pre-computed, although now it is necessary to estimate two parameters: *ζ* and *ξ*. For 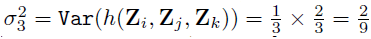 because of symmetry. For large *n* (typically *>* 100), we sub-sample the samples to estimate *α, ζ, ξ* since the sub-sample estimate are reliable and reduce the computation significantly.

**Table 1.** Probabilities required to be estimated for 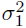 as in eq. (4): (left) *P* (*M* (*i, j, k*), *M* (*i, l, o*)) **only one parameter needed to be estimated (e.g.** *α***). The sum of the rows and columns must be** 1/3 **due to symmetry. (right)** *P* (*m*(*i, j, k*), *m*(*i, l, o*)) **can be pre-computed due to being a scalar assuming ties are improbable.**

|  | i | l | o |  |
| --- | --- | --- | --- | --- |
| i | $\alpha$ | $\beta$ | $\beta$ | $1/3$ |
| j | $\beta$ | $\gamma$ | $\gamma$ | $1/3$ |
| k | $\beta$ | $\gamma$ | $\gamma$ | $1/3$ |
| | $1/3$ | $1/3$ | $1/3$ | |
| i | $2/15$ | $1/10$ | $1/10$ | $1/3$ |
| j | $1/10$ | $7/60$ | $7/60$ | $1/3$ |
| k | $1/10$ | $7/60$ | $7/60$ | $1/3$ |
| | $1/3$ | $1/3$ | $1/3$ | |

**Table 2.** Probabilities required to be estimated for 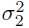. (left) *P* (*MS*(*i, j, k*), *MS*(*i, j, l*)) **only two parameters needed to be estimated (e.g.** *ζ, ξ***). The sum of the rows and columns must be** 1/3 **due to symmetry and i.i.d assumption. (right)** *P* (*m*(*i, j, k*), *m*(*i, j, l*)) **can be pre-computed due to being a scalar assuming ties are improbable.**

|  | i | j | l |  |
| --- | --- | --- | --- | --- |
| i | $\zeta$ | $\eta$ | $\kappa$ | $1/3$ |
| j | $\eta$ | $\zeta$ | $\kappa$ | $1/3$ |
| k | $\kappa$ | $\kappa$ | $\xi$ | $1/3$ |
| | $1/3$ | $1/3$ | $1/3$ | |
| i | $2/12$ | $1/12$ | $1/12$ | $1/3$ |
| j | $1/12$ | $2/12$ | $1/12$ | $1/3$ |
| k | $1/12$ | $1/12$ | $2/12$ | $1/3$ |
| | $1/3$ | $1/3$ | $1/3$ | |

### Remark 5

*For smaller numbers of samples, the variance is better approximated by 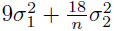or even 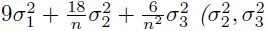 are calculated in the next Remark). In Fig 2 we compare each asymptotic approximation to an exact variance calculated by permutation, to show the rate of convergence.*

**Fig 1.**
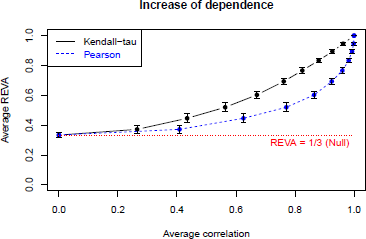
REVA vs Pearson and Kendall correlation in simulated data with controlled correlation between pairwise distances and the scalars. In this case *X*_*i*_’s are scaler i.i.d. with 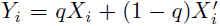 as described in equation (5) The REVA statistic is monotonically increasing with both Kendall-tau and Pearson correlations. As expected, REVA behaves more similarly to Kendall-tau.

**Fig 2.**
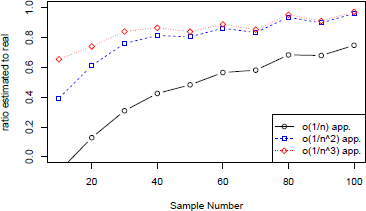
The ratio of asymptotic variance to exact variance, calculated over 1000 permutations data, for different sample sizes. 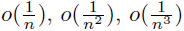 describe increasingly precise approximations to the variance. It can be seen that for small sample numbers, the approximation of 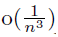 is necessary for accurate approximation but for more samples we can only use 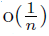. Therefore, the consistent underestimation of the variance is reduced for large sample sizes.

### Remark 6

*A main concern in U-Statistics analysis is the risk of degeneracy of the variance, which occurs when* 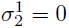 *under the null. Simplifying* 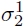 *using the notations in Table 1, and considering the symmetry* 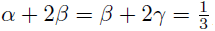, *we can simplify* 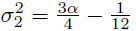. *Hence, we need to show* 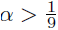. *Using a standard technique, we condition on X*_*i*_, *and use i.i.d. assumption.*

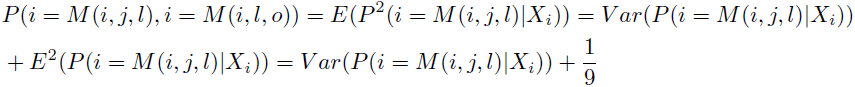

*It follows that as long as Var*(*P* (*i* = *M* (*i, j, l*)*|X*_*i*_)) *>* 0, *or equivalently P* (*i* = *M* (*i, j, l*) |*X*_*i*_) *is not constant in probability, REVA avoids degeneracy. While we cannot entirely rule out the possibility of degeneracy, (we hypothesize that it could happen where the distribution is perfectly symmetric, e.g. in the case of a uniform distribution on a sphere), it is extremely unlikely where there is asymmetry of the distribution or distance measure.*

## Results

### Simulations

To study REVA’s behavior as a correlation measure, we ran a simulation in which we can control the correlation between *X*_*i*_’s and *Y*_*i*_’s. Consider the following one dimensional scenario: Let *X*_*i*_’s be i.i.d standard normals and *D* be L-1 norm. Now, let

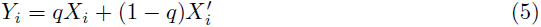

where 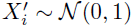 and independent of *X*_*i*_. In this scenario, the Pearson correlation, 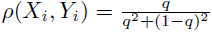. Proposition (1) suggest that if *q* = 0 or *q* = 1, REVA statistic *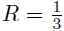* or *R* = 1. To investigate behavior between these extremes, we simulated 1000 rounds under a range of *q* values and show the results as the error bar plot in Fig (1). For Kendall-tau, we used an empirical average for any fixed *q*. As expected, REVA behaves monotonically and more closely resembles Kendall’s coefficient than Pearsons’s. In fact, the following theorem reveals some theoretical similarities:

#### Theorem 2

*Let x and Y be continuous scaler r.v.s. with F*_*XY*_, *F*_*X*_, *F*_*Y*_ *c.d.f.*

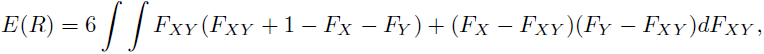

*(while their Kendall’s-τ is* 4 *∫ ∫ F*_*XY*_ *dF*_*XY*_ *-* 1 *[20]).*

#### Proof 2

*Proof of Theorem (2).*

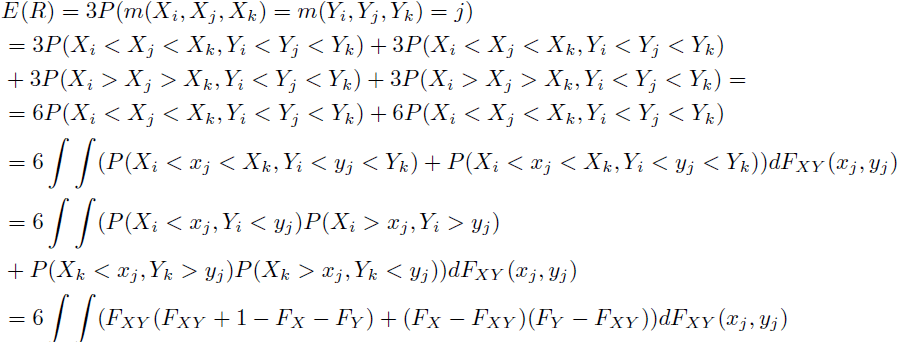

*The first line is because of the symmetry to i, j, k, the second and third line is because all orderings are disjoint, the forth line is due to symmetry i, j, k, the fifth line is due to independence and the sixth line is due to identically distribution of the disjoint of F*_*XY*_.

Expressed in this form, it is easy to see that REVA offers greater power than Kendall in at least one, well-known situation of non-linear association. Consider the example where 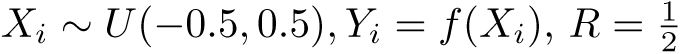 which is bigger than random threshold (i.e 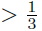.). As known, Kendall-*τ* =0.

One obvious conlcusion of the theorem (2) is the following corollary.

#### Corollary 1

*Under the conditions of theorem (2), REVA > 0.*

#### Proof 3

*Proof of corollary (1): R is non-negative and hence, its expectation, REVA, is non-negative. If E(R) =0, because of positivity of the joint distribution and its complements then*

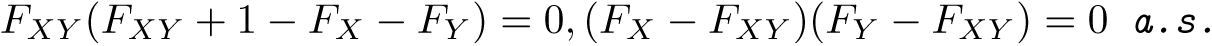

*By subtracting two equality we have F*_*XY*_ *- F*_*X*_*F*_*Y*_ = 0 *a.s. and by replacing back into the second equation, we have F*_*X*_ (1−*F*_*X*_)*F*_*Y*_ (1−*F*_*Y*_) = 0 *almost surely which is contradictory to the definition of the c.d.f of continuous random variables.*

Following Remark (4) and Corollary (1), it makes sense to ask about the minimum value that REVA can assume in the scale case. The following Corollary proves that the example described in Remark 4 achieves the minimum.

#### Corollary 2

*Under the conditions of theorem (2), 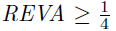*.

#### Proof 4

*Proof of corollary (2): To show that 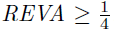, we need to use copula theory to prove the inequality. Applying Sklar’s theorem, REVA can be re-written as*

*F*_*X*_, *F*_*Y*_ *U* (0, 1) *and F*_*XY*_ = *C*(*x, y*) *where C is a copula and x, y ∈* [0, 1]. *Based on the Frechet-Hoeffding bounds* 0, *x* + *y −* 1 *≤ C and C ≤ x, y.*

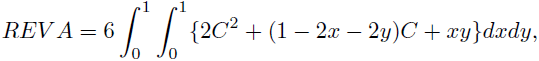

*and since the integrand is positive for any* Ω *∈* [0, 1] *×;* [0, 1],

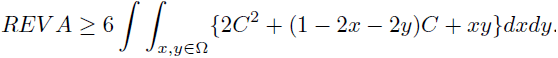

*A specific* Ω, *we are interested in 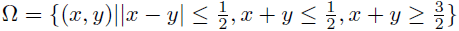. For a fixed x,y, the minimizer of integrand (ignoring the constraint that C is a copula) is when 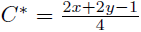. Hence, the integrand 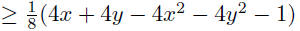. Now, from the last inequality and inequality of the integrand, we have:*

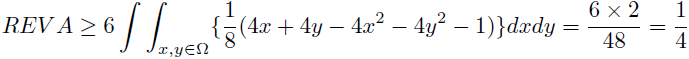

*Note that if* 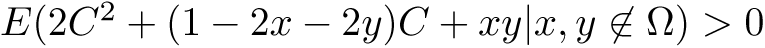, *then 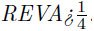. So, a necessary condition for inequality to become an equality is that*

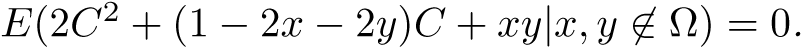

## Robustness Analysis

In this section, we explore the performance of REVA under a variety of simulated conditions. We consider the situation where a simple linear relationship between **X** and *Y* is diminished by increased noise levels, increased dimensionality and the presence of outliers. We also considered a non-linear model, simulating this scenario using the same assumptions about noise, dimensionality and the presence of outliers. In each scenario we compare REVA to the distance correlation approach by Szekely and Rizzo, which has become a standard method for data of this type. Based on the literature of statistics, we expect the rank-based REVA to be less sensitive to influence from outliers, noise, etc. than distance correlation inspired by Pearson’s. Conversely, all else equal, distance correlation should offer better power when the relationship is close to linear. We start with a scenario in which we can control the effects of dimension, noise, etc. Consider the following scenario: Let *Y*_*i*_ *∼ N* (0, 1) and

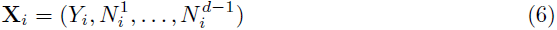

where 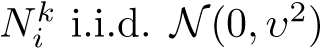 and *D* is the L-1 norm. We vary both parameters *d ∈* {5, 10, 15, 20, 25, 30, 100, 200, 500, 1000, 2000} and *v* ∈ {0.05, 0.10, 0.15, *…,* 0.50}, comparing REVA to Distance Correlation over all combinations of these parameters. The resulting statistics are depicted in the first panels of Fig 3, along with the null case where *X* and *Y* are independent, which is labeled as “Noise Only” in the figure.

**Fig 3.**
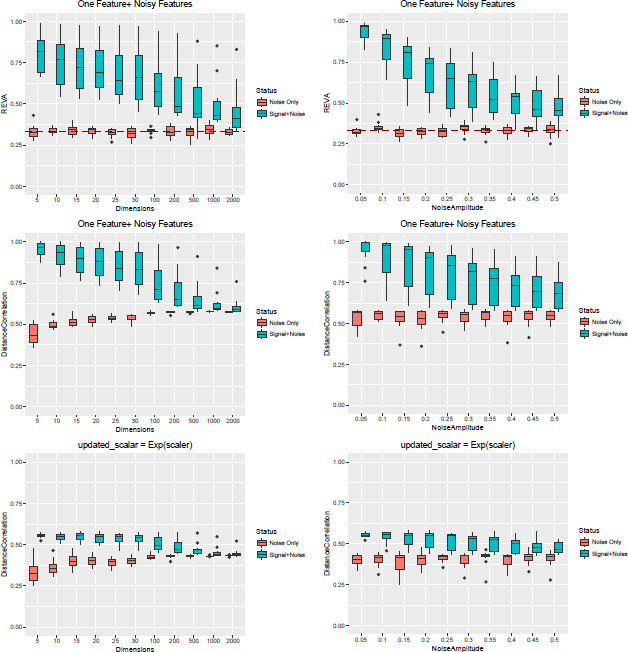
Peformance of REVA (first row) and Distance Correlation (second and third row) under increased dimensionality and/or noise as described above in the “Robustness Analysis” section. In the sub-figures in the top two rows, the “Signal+Noise” scenario is described as in eq. 6 where there is a signal detect and the “Noise only” where there is no signal to detect. The bottom sub-figures depict the distance correlation performance under the scenario in which the scaler has been transformed by a non-linear function, exponential function. Since REVA is ranked-based its outcome is identitical to the first row but distance correlation loses its detection power.

**Fig 4.**
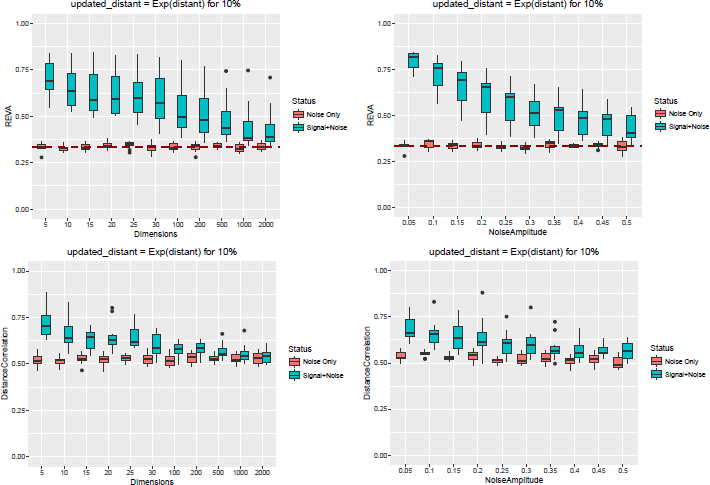
Performance of REVA (top row) and Distance Correlation’s (bottom row) where a randomly selected 10% of pairwise distances were transformed by the exponential function. REVA is more robust to the presence of outliers in the pairwise distances.

**Fig 5.**
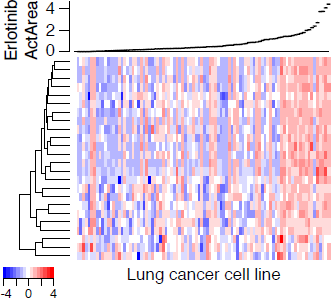
Lung cancer cell lines sorted by response to Erlotinib (Active area, top). Heatmap of gene expression values for corresponding lung cancer cell lines (columns) for the set of epithelial genes colored according to the z-score for each gene (row).

Naturally as the dimension or noise level increases both tests lose power. Distance correlation has better power than REVA in lower dimensions, but as the dimension increases, the relationship is reversed.

We also evaluated performance when the relationship between *X* and *Y* is non-linear, applying an exponential transformation to *Y*, shown in the lower panels of Fig (3). Since REVA is rank-based, it is invariant to monotone transformations, however, distance correlation loses its power dramatically due to violation of the linearity assumption.

We added outliers to the simulation by randomly choosing 10% of the distances and applied the same exponential function used in the first scenario described in this section. The results are depicted in Fig (4). REVA is more robust to outliers than distance correlation. As seen in the scenarios described above, REVA’s distribution under the null is very stable with median very close to 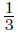. To make a more precise comparison, we calculated the p-value using both REVA and distance correlation in the outlier situation for the “Signal+Noise” scenario and show it in Table 3. Boldface p-values show those under 0.01. As expected, REVA is significantly more powerful distance correlation throughout much of the range of simulated signal to noise ratos.

**Table 3.** Comparison of p-values for REVA’s (top) and Distance Correlation’s (bottom) in simulations performed in Fig (4) for “Signal+Noise.” Columns represent the dimension of X and rows show the noise standard deviation in “noise features.” 10% of pairwise distance were randomly chosen to be manipulated by exponential function. P-values *<* 0.01 are bold-faced. REVA keeps its detection power relatively better than distance correlation as the dimension of feature space or/and the standard deviation of the noise grow, and therefore is more robust to the presence of outliers in the pairwise distances. Due to space limitations some dimensions are dropped.

|  | 5 | 10 | 15 | 20 | 25 | 30 | 100 | 200 | 500 | 1000 | 2000 |
| --- | --- | --- | --- | --- | --- | --- | --- | --- | --- | --- | --- |
| **0.05** | **2e-134** | **1e-127** | **1e-147** | **3e-132** | **2e-125** | **1e-134** | **2e-131** | **3e-111** | **1e-108** | **2e-105** | **1.2e-80** |
| **0.1** | **2e-130** | **1e-110** | **4e-123** | **1e-113** | **2e-110** | **3e-124** | **8e-83** | **9e-73** | **8.2e-56** | **3.9e-38** | **4.9e-25** |
| **0.15** | **7e-103** | **1.3e-97** | **3e-105** | **1e-107** | **2.6e-71** | **7.8e-94** | **1.8e-49** | **2.1e-56** | **1.7e-26** | **1.3e-13** | **4.1e-11** |
| **0.2** | **3e-111** | **4.7e-85** | **1.9e-63** | **2.1e-68** | **3e-61** | **3.3e-57** | **4.1e-30** | **5.4e-32** | **8.9e-11** | **4.3e-07** | **0.0013** |
| **0.25** | **1.9e-75** | **2.6e-66** | **1.5e-46** | **2.2e-50** | **5e-56** | **5.5e-50** | **1.2e-14** | **1.5e-15** | **2.4e-07** | **0.0049** | **4.9e-05** |
| **0.3** | **1.7e-62** | **1.8e-45** | **1.3e-38** | **1.3e-37** | **4e-34** | **5.2e-17** | **4.5e-15** | **1.8e-08** | **2.9e-06** | **0.0079** | **0.0086** |
| **0.35** | **1.5e-52** | **1.7e-38** | **1.5e-23** | **1.3e-21** | **5.6e-24** | **1.1e-27** | **7e-15** | **0.00011** | **3.2e-05** | 0.29 | 0.25 |
| **0.4** | **1.9e-44** | **2.5e-26** | **1.5e-21** | **9.7e-17** | **1.8e-18** | **2.2e-12** | **1.2e-05** | **7e-04** | **4e-04** | 0.03 | 0.085 |
| **0.45** | **6.7e-36** | **2.5e-20** | **2.9e-17** | **5e-17** | **5.4e-22** | **4.1e-11** | **6.2e-05** | **0.00012** | 0.057 | 0.34 | 0.04 |
| **0.5** | **2.2e-26** | **3.4e-22** | **3.6e-16** | **2.2e-20** | **1.8e-09** | **0.00015** | **0.0043** | 0.12 | **0.0022** | 0.035 | 0.59 |
| **0.05** | **2.2e-12** | **6e-10** | **4.8e-05** | **0** | **2.2e-12** | **2e-08** | **3.6e-08** | **1.2e-06** | **1.8e-06** | **2.8e-06** | **0.003** |
| **0.1** | **0.00016** | **0** | **4.8e-05** | **3.4e-08** | **3.2e-11** | **6e-11** | **0.0012** | **0.0015** | **2e-04** | **0.0038** | 0.32 |
| **0.15** | **0.00024** | **0.00063** | **4.2e-11** | **2.2e-16** | **1.9e-09** | **1.1e-09** | 0.013 | **0.00057** | 0.29 | 0.71 | 0.23 |
| **0.2** | **0** | **1.1e-09** | **4.3e-07** | **0.00016** | **9.5e-07** | **4.3e-06** | **0.0077** | 0.042 | 0.051 | 0.65 | 0.26 |
| **0.25** | **3.6e-13** | **6.1e-06** | **4.1e-06** | **0.00011** | **7.1e-05** | **4.6e-05** | 0.059 | 0.017 | 0.13 | 0.55 | 0.38 |
| **0.3** | **0** | **1.1e-05** | **0.00078** | **3.7e-06** | **0.00084** | 0.057 | 0.036 | 0.012 | 0.31 | 0.4 | 0.92 |
| **0.35** | **6.7e-09** | **7.6e-05** | **0.0071** | 0.018 | 0.027 | **0.0075** | 0.037 | 0.45 | 0.29 | 0.7 | 0.15 |
| **0.4** | **4.3e-06** | **0.0041** | **0.00054** | 0.068 | 0.012 | 0.024 | 0.49 | 0.52 | 0.34 | 0.21 | 0.83 |
| **0.45** | **1.5e-05** | 0.05 | 0.041 | **0.00012** | **0.00078** | 0.071 | 0.12 | 0.87 | 0.11 | 0.75 | 0.48 |
| **0.5** | **2.7e-05** | **0.0015** | 0.065 | **0.0041** | 0.041 | 0.73 | 0.52 | 0.7 | 0.6 | 0.06 | 0.42 |

## Application to Genomic Data

An emerging question in cancer research is whether we can predict which patients will respond to a specific therapy based upon the molecular profile of their tumor ([21]). In this analysis, we look at whether lung cancer cell lines with similar gene expression profiles will respond similarly to Erlotinib, a therapeutic agent approved by the FDA for use in lung cancer. The data for this analysis is obtained from the Cancer Cell Line Encyclopedia (CCLE, [22]). Genome-wide gene expression values for hundreds of cancer lines were obtained using the Affymetrix hgu133plus2 arrays. Drug response for each of these cell lines is reported as ActiveArea, a summary of the rate at which cancer cells are killed across a range of dose levels, such that larger values indicate greater sensitivity to the drug.

We chose this example because the mechanism of action for this drug is well understood, permitting us to make predictions about the outcome of the study. Specifically, erlotinib inhibits the Epidermal Growth Factor Receptor (EGFR) gene, which is a commonly activated and serves as an oncogene in lung cancer. Therapeutic response to EGFR inhibition has been associated with a biological processes called the epithelial to messenchymal transition (EMT) in numerous cancer types [23–25]. Therefore, we would expect genes associated with that pathway and response to erlotinib to follow the neighbors principle. Prior studies [26] have defined a robust EMT signature in lung cancer which can be segregated into two sets of genes, *epithelial* and *mesenchymal* genes. We apply REVA separately to each set, to test whether the neighbors principal holds in between erlotinib response and gene expression profiles of either the epithelial or messenchymal genes using L-1 distance. We confirm that REVA finds that gene expression profiles for epithelial genes are more significantly associated with Erlotinb response (p-value of 3.6e-08 for R = 0.392, 5) than messenchymal genes (p-value of 0.033 for R=0.356, not shown).

## Conclusion

In this paper, we seek a robust, generalized measure of correlation for the case where at least one variable is multidimensional. Although several relevant methods have been developed in recent years [5–9], there is only one rank-based test available to date [9], and it violates the principle that correlations should be symmetric in the samples. We introduce REVA which extends Kendall’s rank-based *τ* statistic to operate on pairwise point distances for triplets of points. The result is nonparametric and rank based, and symmetrical in the samples. It is very flexible, capable of capturing an array of non-linear relationships among the variables in addition to linear relationships. The expected value of REVA under the null hypothesis is constant (equal to 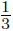), it does not depend on the dimensionality of the data and is asymptotically normal. Therefore, REVA can be quickly and reliably computed for datasets with large feature spaces. The resulting statistics are slightly more computationally demanding than the alternative methods. REVA is *O*(*n*^3^) as compared to *O*(*n*^2^ log *n*) for Heller’s and Shen’s approach and *O*(*n*^2^) for distance correlation, where *n* is the sample size. Nonetheless, although a thorough analysis of the performance of REVA is beyond the scope of this paper, our results on simulated data in comparison to distance correlation suggest that REVA is a powerful test in some situations. Work is ongoing on extensions of the REVA approach. Shen et al. extended distance correlation to capture a wide array of non-linear relationships by calculating a local correlation, rather than weighing all pairwise relationships among variables equally [9] The same approach can be implemented with REVA.

An interesting inverse question arises from work in cancer genomics. Recent data suggests that more aggressive tumors have more heterogeneous genetic profiles ([2, 17, 27]). In this case, we would expect a stronger relationship between tumor profile and response in indolent tumors than in their rapidly growing counterparts. The goal of this analysis is then a change-point problem, where the goal is to identify points along the *Y* scale where the correlation between **X** and *Y* changes. Similar approaches have been applied for time course analysis in genomics [28]. We believe that the REVA framework is amenable to reformulation, providing a general framework for non-parametric, rank-based change point analysis applicable both for prediction and time course analysis.

## Acknowledgments

This work was supported by the National Institutes of Health (grant P30 CA006973) ant Russian Science Foundation (grant 17-00-00208 KOMFI)

